# Predicting cognitive abilities across individuals using sparse EEG connectivity

**DOI:** 10.1101/2020.07.22.216705

**Authors:** Nicole Hakim, Edward Awh, Edward K Vogel, Monica D Rosenberg

**Affiliations:** Department of Psychology, University of Chicago, Chicago, Il; Institute for Mind and Biology, University of Chicago, Chicago, Il; Grossman Institute for Neuroscience, Quantitative Biology, and Human Behavior, University of Chicago, Chicago, I

## Abstract

Human brains share a broadly similar functional organization with consequential individual variation. This duality in brain function has primarily been observed when using techniques that consider the spatial organization of the brain, such as MRI. Here, we ask whether these common and unique signals of cognition are also present in temporally sensitive, but spatially insensitive, neural signals. To address this question, we compiled EEG data from individuals performing multiple working memory tasks at two different data-collection sites (*ns* = 171 and 165). Results revealed that EEG connectivity patterns were stable within individuals and unique across individuals. Furthermore, models based on these connectivity patterns generalized across datasets to predict participants’ working memory capacity and general fluid intelligence. Thus, EEG connectivity provides a signature of working memory and fluid intelligence in humans and a new framework for characterizing individual differences in cognitive abilities.

## INTRODUCTION

Human brains share a common template of functional organization. Nearly every person, for example, shows a retinotopic map in primary visual cortex (Engel et al., 1997) and a face-sensitive region in inferior temporal cortex (Kanwisher et al., 1997). Electroencephalogram (EEG) activity signatures of the number of items in working memory are reliable enough to be detected at the single-subject level (Vogel & Machizawa, 2004). Even in the absence of an explicit task, individuals show synchronous activity in a stereotyped set of brain networks, such as the default mode (Damoiseaux et al., 2006) and frontoparietal (Duncan & Owen, 2000) networks.

Atop this shared organizational template is significant variability in functional brain architecture. Functional MRI studies have revealed that each person has a unique pattern of synchronous activity between spatially distinct brain regions, known as functional connectivity, that distinguishes them from others and remains stable across cognitive states (Finn et al., 2015; Gratton et al., 2018; Miranda-Dominguez et al., 2014). Furthermore, these unique patterns, or whole-brain connectomes, appear cognitively meaningful, predicting individual differences in behaviors including fluid intelligence (Finn et al., 2015; Greene et al., 2018) and attention (Kessler et al., 2016; O’Halloran et al., 2018; Poole et al., 2016; Rosenberg et al., 2016).

Brain-based predictive models rely on this interesting duality in brain function: broadly similar organization with consequential individual variation. That is, a neural system common across individuals is necessary to build brain-based biomarkers because if every individual relied on a different neural system to achieve a particular behavior, predictive models would fail to generalize across people. However, systematic idiosyncrasies in these common brain networks are what allow models to predict each person’s unique set of cognitive abilities. Without differences in these common neural systems, predictive models would fail to differentiate individuals. This duality in brain function has primarily been observed when using techniques, such as MRI, that consider the spatial organization of the brain. In this paper, we seek to address the open theoretical question of whether these common and unique signals of cognition are also present in temporally sensitive, but spatially insensitive, neural signals, such as EEG.

Based in part on evidence from connectome-based models of behavior, work in cognitive and network neuroscience has argued that cognition relies on coordinated activity in large-scale, high-density brain networks (Bressler & Menon, 2010; Medaglia et al., 2015; Park & Friston, 2013). This may suggest that dense functional networks revealed by techniques with relatively high spatial resolution, such as fMRI, are uniquely informative of behavior. This hypothesis has not been directly tested. However, studies relating cognitive abilities to brain connectivity patterns measured with electrical or magnetic signals (i.e., using electroencephalography [EEG] or magnetoencephalography [MEG]; Burgess & Ali, 2002; Damaševičius et al., 2018; Fellrath et al., 2016; Karamzadeh et al., 2013; Nentwich et al., 2020; Palva et al., 2010)—which often include fewer than one percent of the spatial resolution typically available to fMRI functional connectivity models—suggest that cognitively meaningful variability in brain function may be reflected in sparse, high-frequency neural signals. In this paper, we directly test the hypothesis that dense functional networks are uniquely informative of behavior. To do so, we ask whether sparse EEG connectivity can 1) identify individuals and 2) predict trait-like cognitive abilities across individuals from completely independent datasets.

In addition to open theoretical questions about brain function and cognition, obstacles to widespread adoption of existing connectome-based models of behavior remain. First, the majority of MRI-based models have not tested the generalizability of their predictions to unseen individual and datasets (Woo et al., 2017). This limits our ability to draw conclusions about their robustness and replicability (Poldrack et al., 2020). Second, work has suggested that, in some cases, confounds including head motion can influence observed relationships between fMRI networks and behavior (Siegel et al., 2017). Finally, the costs of MRI for researchers, clinicians, and participants has so far limited translation to real-world settings.

Here we address these open theoretical questions and practical challenges by using a direct measure of neural activity that is easy and affordable to implement: EEG. Across two EEG datasets, each with 165+ individuals, collected at different universities with different EEG systems (passive vs. active), we show that trial-evoked EEG functional connectivity patterns are unique across individuals and stable within individuals. We next demonstrate that models based on sparse EEG connectivity patterns generalize across individuals and independent datasets to predict individual differences in working memory capacity, a critical cognitive ability. Finally, we show that the same EEG connections that predict working memory capacity predict general fluid intelligence in novel individuals. Thus, sparse EEG connectivity patterns reveal a signature of trait-like cognitive abilities in humans and provide a new, more affordable and accessible approach for predicting cognitive ability from brain function.

## RESULTS

### Whole-scalp evoked EEG connectivity fingerprinting

Are patterns of EEG activity unique across individuals? Many EEG analyses implicitly treat potential individual differences as noise, averaging results across the group or comparing average results from two groups (e.g., individuals with high vs. low working memory capacity; patients vs. controls). Here, we asked whether individuals have both *stable* and *unique* trial-evoked EEG connectivity patterns that can reliability distinguish them from others. To test this possibility, we applied “functional connectivity fingerprinting”, an approach developed using fMRI functional connectivity data (Finn et al., 2015), to two independent EEG datasets. Data were collected as participants performed variants of a lateralized change detection task at the University of Oregon (*n* = 171) and the University of Chicago (*n* = 165; see Methods for details). Fingerprinting analyses were restricted to individuals who participated in at least two separate tasks at one of the sites (Oregon *n*=171; Chicago *n*=45).

For each participant and task, we calculated a whole-scalp evoked EEG connectivity pattern using the 17 electrodes that overlapped between the two datasets. We correlated the trial-averaged time courses between all pairs of these 17 electrodes for each task separately. In other words, we operationalized EEG connectivity as the temporal correlation between event-related potentials (ERPs) in two electrodes. This analysis generated at least two EEG connectivity matrices for each individual. Next, we vectorized the matrices for each individual and task. Finally, we correlated every individual’s task *A* matrix with all possible task *B* matrices and vice versa. Individuals were considered correctly identified when the correlation between their task *A* and their task *B* connectivity patterns was larger than the correlation between their task *A* and all other task *B* connectivity patterns (and vice versa). Identification accuracy is the number of correct identifications divided by twice the number of participants. Non-parametric statistical significance was determined with 10,000 permutation tests (see Methods for detail). As a comparison, we also characterized the stability and uniqueness of EEG amplitude (see Supplemental Materials for more detail).

We additionally ran an even stronger test of individual uniqueness by directly comparing the effects of subject vs. task. To do this, we correlated each individual’s task *A* and task *B* matrices (“across-task correlations”). We then correlated each individual’s task *A* matrix with everyone else’s task *A* matrix (“within-task correlations”). We also calculated these within-task correlations for task *B*. We then compared the across-task and within-task correlations. If individuals look most like themselves, regardless of task context, then across-task correlations should be higher than within-task correlations.

#### University of Oregon sample

Participants in the University of Oregon sample completed two lateralized change detection tasks in which they were instructed to remember items’ shape or color as part of a single study. Functional connectivity fingerprinting analyses revealed that individuals had stable and unique evoked EEG connectivity patterns across these two tasks that could reliably dissociate them from other individuals: color: shape task accuracy = 82%, p<0.0001; shape: color task accuracy = 81%, p<0.0001 (chance = 1/171, or .58%; **Figure 1**). In other words, individuals’ EEG connectivity patterns were distinct from the group and stable across tasks. Furthermore, individuals’ EEG connectivity networks looked most like themselves, regardless of task context (across-task matrices average=0.9992+/-0.00058; within-task matrices average=0.9941+/-0.0039; p<0.0001).

**Figure 1.**
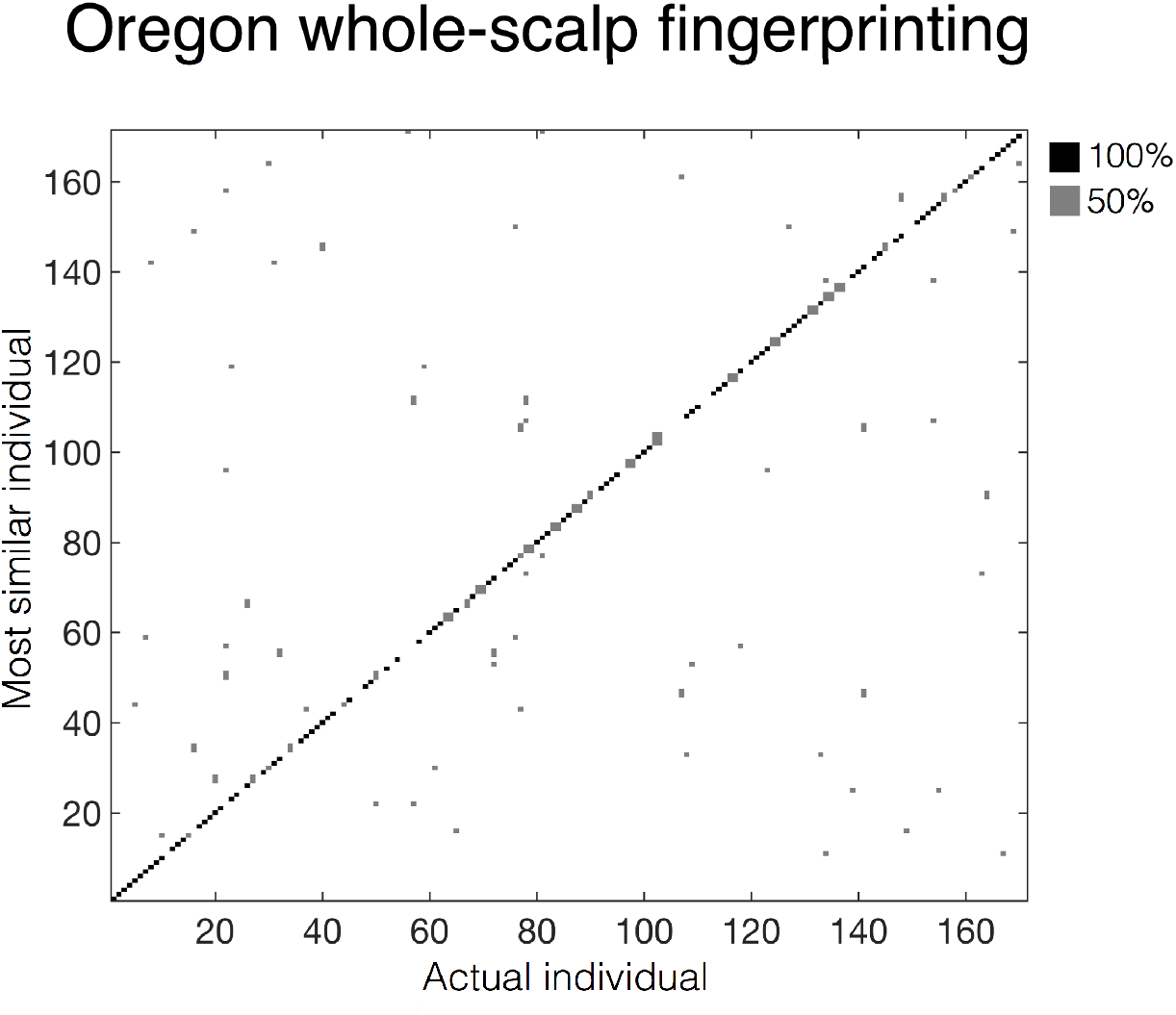
Functional connectivity fingerprinting results from Oregon-site data. The x-axis represents the actual individual, whereas the y-axis represents the most similar individual. Black squares indicate that an individual was most similar to the same person in both the color and shape tasks. Gray squares indicate that an individual was identified as a different people in the color and shape tasks. Both axes are organized based on working memory capacity, with lower working memory capacity individuals represented by lower numbers and vice versa. Any square along the diagonal indicates that an individual was correctly identified.

#### University of Chicago sample

The full University of Chicago sample included data from twelve experiments collected as part of six independent studies. Data from all studies were collected on different days in different sessions. For this analysis, we only consider data from the subset of individuals who participated in multiple experiments (see Methods for detail). Replicating results from the Oregon sample, individuals in the Chicago dataset showed stable and unique evoked EEG connectivity patterns, which could be used to reliably distinguish them from other individuals: task *A:B accuracy=29% p*<0.0001; task *B:A accuracy=34%, p*<0.0001 (chance = 1/19, or 5.26%). Additionally, individuals’ EEG connectivity network looked most like themselves, regardless of task context (across-task matrices average=0.9955+/-0.0046; withintask matrices average=0.9923+/-0.0063; p<0.0001).

Although identification accuracy was numerically lower in the Chicago than the Oregon dataset, this was expected as the Chicago analyses identified individuals viewing distinct displays from different testing sessions, whereas the Oregon analyses identified individuals across tasks within the same testing session. Thus, these results demonstrate that individuals’ EEG connectivity patterns remained stable even across different days and modest variations in electrode position.

In both the Chicago and Oregon datasets, we identified individuals based on their unique pattern of EEG connectivity. These results suggest that individuals have unique and robust patterns of evoked EEG connectivity that differentiate them from others. To test whether individual differences in these patterns reflect individual differences in cognitive abilities and behavior, we next asked whether we could use evoked EEG connectivity to predict a central cognitive ability: working memory capacity.

### Predicting working memory from evoked EEG connectivity: Within-site validation

To determine whether EEG connectivity measured during lateralized change detection tasks predicted working memory capacity in novel individuals, we trained and tested connectome-based predictive models (Rosenberg et al., 2016; Shen et al., 2017) using balanced 5-fold crossvalidation. For each participant, we calculated working memory capacity (K score) based on their change detection task performance (Cowan, 2001; Pashler, 1988). We calculated evoked EEG connectivity for each participant as described above. However, instead of separating the data by task, we averaged trials from all experiments and then calculated one EEG connectivity matrix per participant. In each cross-validation fold, we identified the edges that were significantly related to K score in the training sample and used these edges to calculate a summary feature of EEG network strength for each participant (see Methods for more details on edge selection). Using a linear model, we then related these EEG connectivity summary features to K scores. Finally, we used this model to predict the left-out set of participants’ K scores based on the strength of each of their EEG networks. To assess predictive power, we calculated the Pearson correlation between observed and predicted K scores. Corresponding *p*-values were calculated using non-parametric permutation tests. As a comparison, we also predicted working memory capacity using amplitude, rather than EEG connectivity (see Supplemental Materials for more detail).

#### University of Oregon sample

We averaged each participant’s evoked EEG connectivity pattern from the color and the shape change detection tasks and built models to predict working memory capacity measured in a completely independent task. Models significantly predicted novel individuals’ working memory capacity: median correlation between predicted and observed K score *r*=0.25,*p*<0.0085, *mse=0.24, Cohen’s d=2.58;* **Figure 2**.

**Figure 2.**
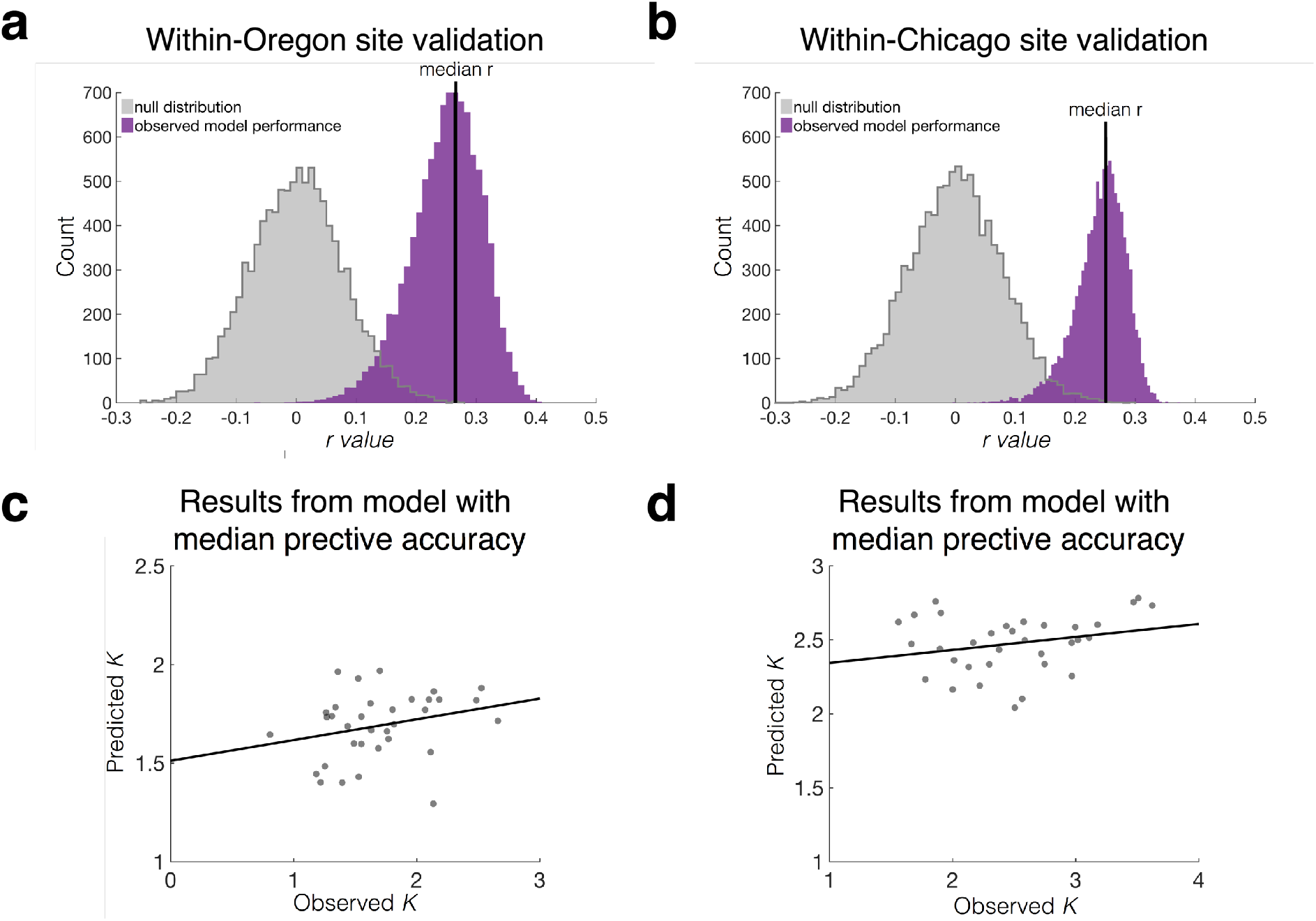
Within-site validation results. Histogram of the correlation between observed and predicted working memory capacity (*K*) using 5-fold cross-validation over 10,000 true model iterations (purple) and 10,000 null model permutations (gray) for (**a**) the model trained within the Oregon-site data and (**b**) the model trained within the Chicago-site data. The vertical black lines represent the iteration and fold with the median *r* value. Scatter plot of the correlation between observed and predicted K scores from the iteration and fold with the median *r* value for (**c**) the model trained within the Oregon-site data and (**d**) the model trained within the Chicago-site data. Gray dots represent individuals observed and predicted K scores from one fold. The black line is the best fit line.

Next, we built predictive models for the color and shape tasks separately. We found that observed and predicted K scores were significantly correlated in the shape task, but not the color task (median shape task, *r*=0.26,*p*<0.0045, *mse*=0.42, *Cohen’s d*=3.00; color task, *r*=0.13, *p*<0.08, *mse*=0.25, *Cohen’s d*=1.38).

We next investigated whether models generalized across tasks by applying models trained in the color task to shape task data from held-out individuals and vice versa. Models trained on color task data significantly predicted K scores from shape-task connectivity (median correlation between predicted and observed K score *r*=0.26, *p*<0.0045, *mse*=0.45, *Cohen’s d*=2.74). Models trained on data from the shape task, however, did not significantly predict K scores from color-task connectivity (*r*=0.13,*p*<0.08, *mse*=0.42, *Cohen’s d*=1.38).

#### University of Chicago sample

For the Chicago dataset, we collapsed data across multiple lateralized change detection experiments, which had different memoranda and sample durations. We calculated K for each participant from the EEG task data. We once again applied 5-fold cross validation to determine whether EEG connectivity could predict working memory capacity across individuals. Replicating findings from the Oregon sample, observed and predicted K scores were significantly correlated: median *r*=0.24,*p*<0.0044, *mse*=0.33, *Cohen’s d=2.71,* **Figure 2**. Thus, evoked EEG connectivity significantly predicts working memory capacity across individuals.

### Predicting working memory from EEG connectivity: Across-site validation

Although these results provide the first evidence that models based on EEG data predict working memory capacity across individuals, there are many reasons that internal (i.e., within-dataset) validation may overestimate effect sizes, including idiosyncrasies in task context, EEG systems, and participant populations. Therefore, an even more powerful demonstration of the robustness of predictive models is to externally validate them—that is, to apply them to data from a completely independent sample. External validation allows us to better approximate a model’s population-level generalizability and better understand its predictive boundaries.

To test the cross-dataset generalizability of models predicting working memory capacity, we trained models on the full Oregon sample and applied them to the full Chicago sample and vice versa. Importantly, these two datasets were collected by different experimenters in different locations using different EEG systems.

#### Predicting Chicago K scores from Oregon connectivity model

To generalize the Oregon working memory network model to the Chicago dataset, we trained a model using the Oregon dataset. To select the most reliably predictive edges from the Oregon data, we calculated the top 5% of positive and top 5% of negative edges (10% of total edges) in the color and shape task separately. We defined the predictive edges as those edges that were included in the top 10% of both tasks. Next, we calculated EEG connectivity summary features for all 171 individuals in the Oregon dataset using these predictive edges. We then related these summary features to K scores in the Oregon dataset using a linear model. Finally, using the same predictive edges, we calculated summary features for each of the 165 individuals from the Chicago dataset. We input these summary features into the Oregon model to generate a predicted K score for each individual in the Chicago dataset. Predictions from this model were significantly correlated with observed K scores, demonstrating that the model trained on the Oregon dataset significantly predicted working memory capacity in the Chicago dataset: *r*=0.24, *p*=0.002, *mse*=0.95; **Figure 3**.

**Figure 3.**
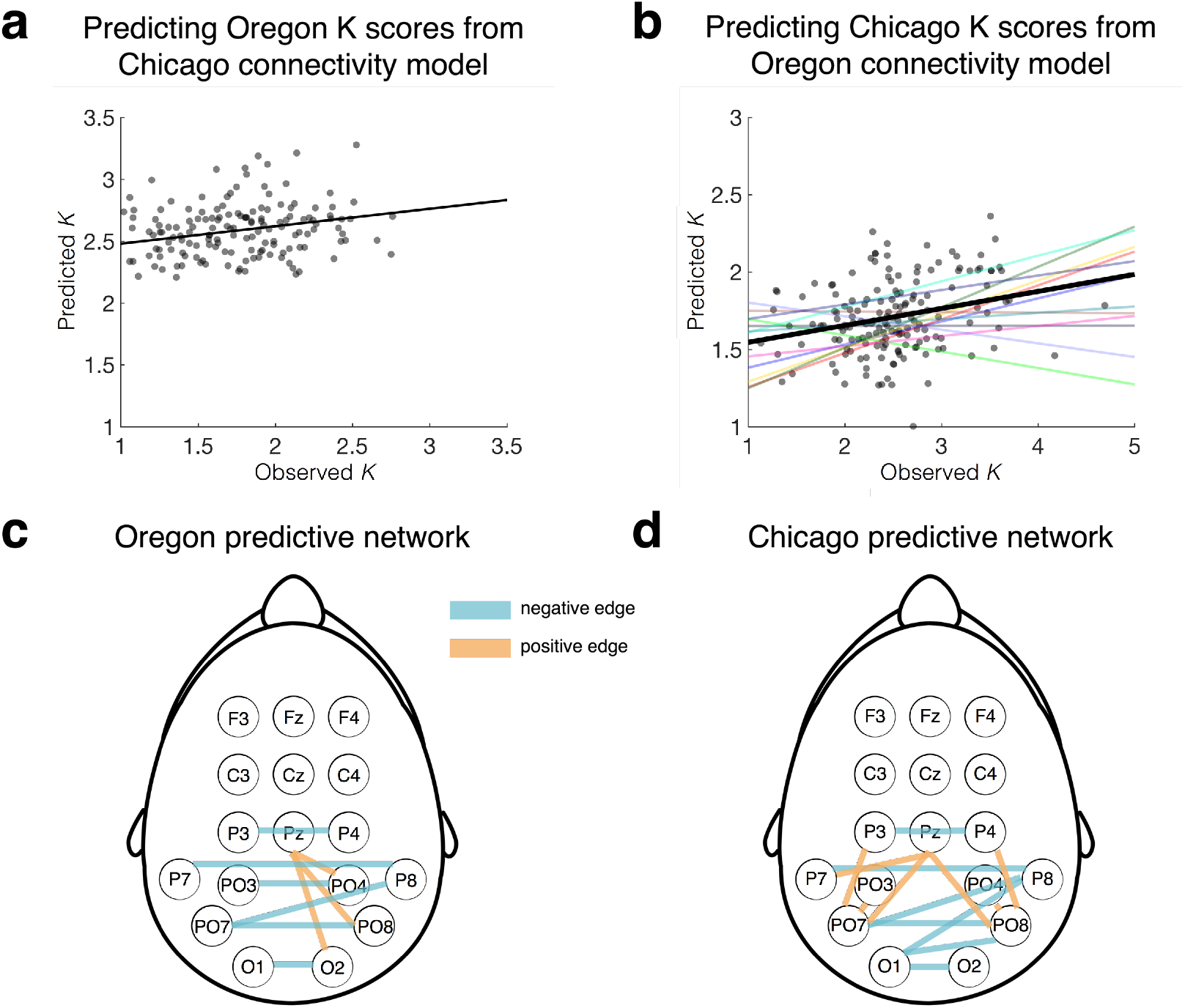
Across-site validation results. **a** Scatter plot of the correlation of observed and predicted working memory capacity (*K*) from a model trained on all of the Chicago data and tested on all of the Oregon data. Gray dots represent individuals observed and predicted K scores. The black line is the best fit line. **b** Scatter plot of the correlation of observed and predicted working memory capacity (*K*) from a model trained on all of the Oregon data and tested on all of the Chicago data. Individual colored lines represent the correlation between observed and predicted working memory capacity of each experiment plotted separately, and demonstrate that it is not the case that a small subset of experiments drives these results. **c** Predictive edges that were included in the model trained on the Oregon dataset. Edges that positively predicted working memory capacity are depicted in orange, and those edges that negatively predicted working memory capacity are depicted in blue. **d** Predictive edges that were included in the model trained on the Chicago dataset.

#### Predicting Oregon K scores from Chicago connectivity model

We next applied a model defined using Chicago data to predict working memory in the Oregon sample. The predictive network was defined as the top 5% of positive and top 5% of negative predictive edges from a model trained on all of the Chicago data. We elected to take this approach rather than taking the overlap of models defined on each task separately because each task had a relatively small sample size (*n*s = 19–29), which could result in unreliable features. Using the predictive edges from the Chicago sample, we then calculated the EEG connectivity summary features for each of the 171 individuals in the Oregon dataset, and input these values into the linear model defined using the full Chicago sample. Predicted and observed K scores were significantly correlated, demonstrating that the model trained on the Chicago dataset significantly predicted working memory performance in the Oregon dataset: *r*=0.28, *p*=0.0003, *mse*=1.06; **Figure 3**.

With this across-site validation, we demonstrate that models predicting working memory capacity not only generalized across EEG session and stimuli, but also generalized across data acquisition sites (Oregon vs. Chicago) and EEG systems (passive vs. active). This suggests that EEG connectivity is a robust, reproducible, and generalizable predictor of working memory capacity.

### Predictive network anatomy

#### Overlap of Oregon and Chicago predictive networks

To determine whether the two models predicted working memory from a common underlying brain network, we tested whether the predictive edges in the Oregon network (9 edges) and Chicago network (14 edges) significantly overlapped using a hypergeometric cumulative density function (e.g., Rosenberg et al., 2016). We found that indeed there was significant overlap between the predictive edges in the Chicago and Oregon networks: 6 overlapping edges (26% of the edges in both networks; 5 edges negatively predicting behavior in both samples and one edge positively predicting behavior in both samples), *p*=7.68e-7. This suggests that a common network predicts working memory capacity in both samples.

#### Network anatomy

The edges that overlapped between the Oregon and Chicago networks involved posterior and occipital electrodes: O1-O2, PO8-PO7, P7-P8, PO7-P8, P4-P3, PO8-Pz. Most of these edges were cross-hemisphere connections, which could be due to the lateralized nature of the task. Interestingly, this predictive network includes electrodes that are typically used to calculate the contralateral delay activity (CDA), an EEG signal that tracks the amount of information held in working memory.

To determine whether the observed working memory capacity network was simply tracking the CDA, we trained models to predict working memory capacity using only the CDA. To do this, we calculated the CDA by taking the difference in amplitude between contralateral and ipsilateral posterior/occipital electrodes (P7/P8, P3/P4, PO7/PO8, PO3/PO4, O1/O2) from 0-1000 ms (the time range included in our EEG connectivity analyses). We then trained a linear model to predict working memory capacity from the CDA using the full Oregon-site dataset and tested it on the full Chicago-site dataset (and vice versa). These models did not significantly predict working memory capacity (train Oregon, test Chicago: *r*=0.04, *p*=0.61; Train Chicago, test Oregon: *r*=-0.32, p=1.74e-5). The significant negative correlation does not provide evidence that the model trained on the Chicago-site data generalizes to predict working memory capacity in the Oregon-site data. This model predicted K score values that ranged from 2.50 to 2.52. This lack of variability in the predictions caused the correlation to be significant, even though the actual predictions are not meaningful. This suggests that, despite the overlap in CDA electrodes and the predictive edges, the connectivity models appear to rely on neural signals that are distinct from the CDA.

To further investigate whether the CDA explains unique variance to EEG connectivity, we trained models to predict working memory capacity from both CDA and evoked EEG connectivity. These models significantly predicted behavior (train Oregon, test Chicago: *r*=0.25, *p*=0.002; train Chicago, test Oregon: *r*=0.27, *p=*0.0004). However, the CDA intercept was only significant in the model trained on the Oregon data (train Oregon CDA intercept: p=0.02; train Chicago CDA intercept: p=0.61). These results provide further evidence that our evoked EEG connectivity models do not simply track the CDA. Instead, they predict behavior from neural signals that are unique.

#### EEG connectivity fingerprinting using predictive networks

Are evoked EEG connectivity networks that predict working memory capacity across individuals reliable enough to distinguish individuals from a group? To ask this question, we applied the same functional connectivity fingerprinting analysis described in the *Whole-scalp EEG connectivity fingerprinting* section above. However, instead of using the whole-scalp network, we only included the edges that significantly predicted behavior across individuals. This network only included those edges that significantly predicted behavior in both the Chicago and Oregon models. We then compared the results from the predictive network to those from the whole-scalp network.

##### University of Oregon sample

Edges predicting working memory were sufficient to significantly identify individuals: color: shape task accuracy=40%, *p*<0.0001; shape: color task accuracy=39%, *p*<0.0001 (chance = 1/171; **Figure 1**). In other words, the EEG connectivity network that predicted working memory capacity across individuals also reliably distinguished individuals. However, identification accuracy using only predictive edges was lower than when we included the whole-scalp network. To determine whether this reduction in identification accuracy was due to down-sampling the number of edges, we compared identification accuracy of predictive edges to an equal number of randomly selected edges that were not predictive of working memory performance. Identification accuracy using a random subset of edges also significantly identified individuals: color: shape task accuracy=39%, *p*<0.0001; shape: color task accuracy=33%, *p*<0.0001 (chance = 1/171; **Figure 1**). There was not a significant difference between identification accuracy using the predictive edges and the random edges: color: shape *p*<0.167; shape: color *p*<0.272. These results suggest that edges that significantly predict working memory capacity are not necessarily better at identifying individuals than a random subset of edges.

##### University of Chicago sample

Replicating results from the Oregon sample, identification accuracy using only predictive edges was successful: task *A*:*B* accuracy=26%, *p*<0.0001; task *B:A* accuracy=23%, *p*<0.0001 (chance = 1/19). This suggests that the network that predicted working memory capacity across individuals was also stable across tasks and was able to distinguish individuals from a group. Once again, identification accuracy using only predictive edges was worse than when we included the whole-scalp network. To determine if this decrease in identification accuracy was due to down-sampling the number of edges, we calculated identification accuracy of a random subset of edges. We found that these random edges also significantly identified individuals: task *A:B accuracy*=13%, *p*<0.0001; *B:A accuracy*=13% *p*<0.0001 (chance = 1/19). However, unlike in the Oregon dataset, the predictive edges more accurately identified individuals than the random edges: task *A*:*B p*<0.0001; task *B*:*A p*<0.0001. These results suggest that edges that significantly predict working memory capacity may more accurately identify individuals than a random subset of edges. Given the inconsistency of this result across the Chicago and Oregon samples, further research is needed to characterize whether an individual’s most identifiable edges are the same edges that best predict their behavior.

### Relationship between evoked EEG connectivity and other cognitive abilities

Our results demonstrate that EEG connectivity observed during a working memory task is a robust and reliable predictor of working memory capacity. Does EEG connectivity observed in this context predict other cognitive abilities? In the Oregon dataset, participants performed a series of cognitive tasks outside the EEG booth (see Methods section for details). Therefore, we used this dataset to investigate the relationship between EEG connectivity and other cognitive abilities.

#### Predicting fluid intelligence from EEG connectivity

We first tested whether networks that predict working memory capacity also predict general fluid intelligence (gF), a closely related ability (Conway et al., 2002; Engle et al., 1999; Fukuda et al., 2010). To this end, we calculated gF scores for each participant in the Oregon sample from a combination of performance on three tasks: Raven advanced progressive matrices, number series, and Cattell’s culture fair test, as reported in previous work (Unsworth et al., 2015).

To determine whether the same EEG connectivity networks that predict working memory capacity also predict gF in novel individuals, for each subject in the Oregon dataset, we calculated the strength in the working memory network defined in the Chicago sample. Next, using 5-fold cross-validation, we defined a linear model relating these working memory network strength values to gF. We then used this model to predict the left-out set of participants’ gF scores. Models significantly predicted novel individuals’ gF scores: *r*=0.29, *p*<7.00e-4, *mse*=6.87, *Cohen’s d*=3.09; **Figure 4**. These results suggest that EEG connectivity is not specific to measures of working memory capacity. Although the EEG data used here were collected as participants performed a working memory task, EEG connectivity generalized to predict fluid intelligence. Of note, this modeling approach is similar to correlating predicted K and observed gF, except that the linear model has a gF-specific coefficient that changes in every fold. Both observed and predicted K scores are correlated with gF (observed K: *r*=0.41, *p*=5.98e-7; predicted K: *r*=.18, *p*=.038), suggesting that retaining the model is not critical.

**Figure 4.**
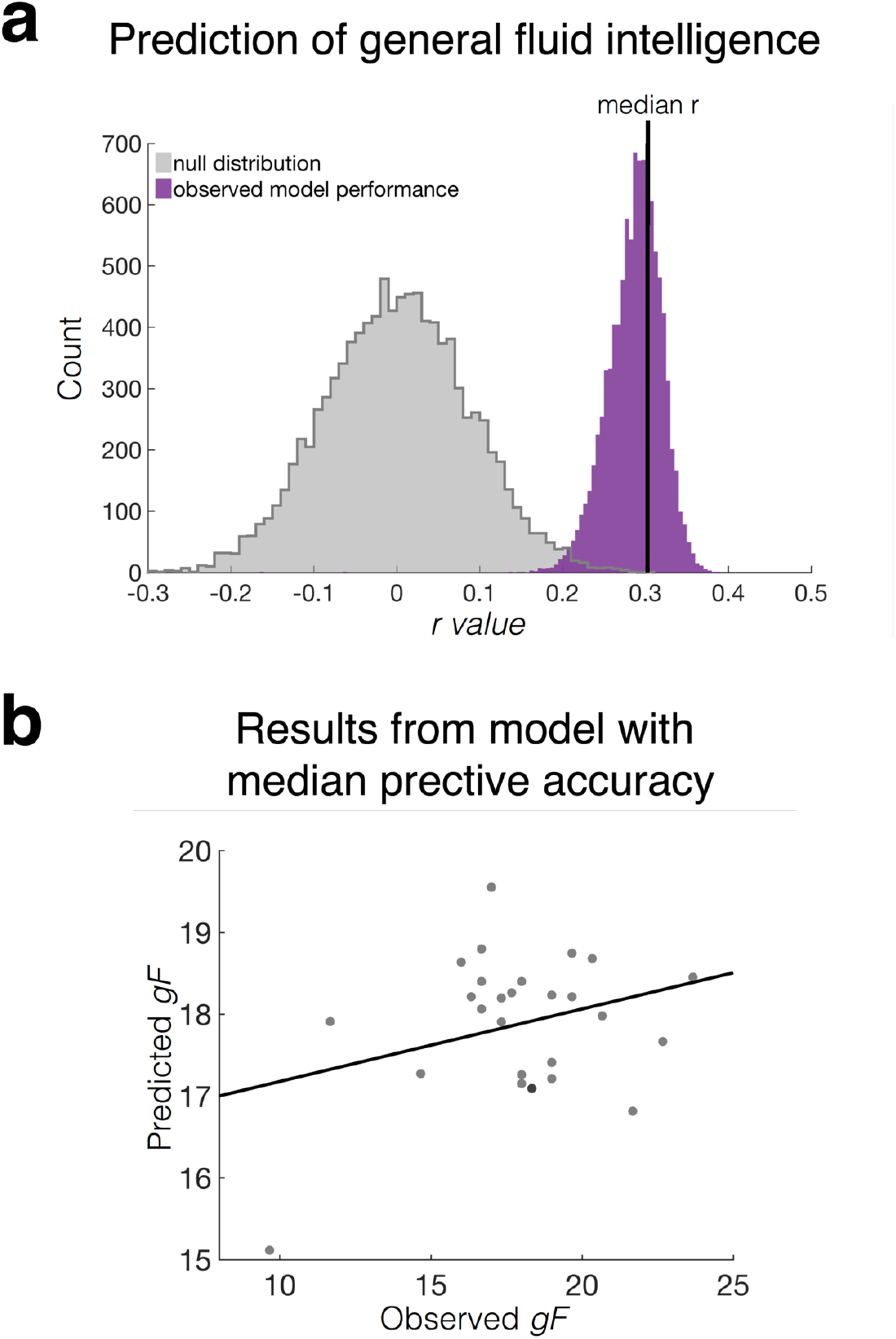
Prediction of general fluid intelligence. **a** Histogram of the correlation between observed and predicted general fluid intelligence (*gF*) using 5-fold cross-validation over 10,000 true model iterations (purple) and 10,000 null model permutations (gray). The vertical black line represents the iteration and fold with the median *r* value. Models were trained and tested within the Oregon dataset using the predictive working memory edges that were included in both the Oregon and Chicago working memory models. **b** Scatter plot of the correlation between observed *gF* and predicted *gF* from the iteration and fold with the median *r* and *p* values. Gray dots represent individuals observed and predicted *gF.* The black line is the best fit line.

#### Correlations between predicted working memory capacity and other cognitive abilities

We next characterized how predicted working memory capacity from our models related to fluid intelligence as well as other cognitive abilities measured in the Oregon dataset, including attentional control and long-term memory. To investigate this, we performed a behavioral crosscorrelation analysis. First, we correlated observed K scores with all other observed behavioral scores (**Table 1**, column 2). Next, we correlated predicted K scores and all other observed behavioral scores (**Table 1**, column 3). Correlating these two columns of data revealed that the relationship between *predicted* working memory capacity and other cognitive abilities mirrors the relationship between *observed* working memory capacity and other cognitive abilities: *r*=0.90, *p*=7.17e-10. In other words, model predictions capture variance in working memory that is shared with related cognitive abilities, rather than variance that is specific to the working memory task used in the EEG studies.

**Table 1.** Behavioral cross-correlation. Characterization of the performance on the additional behavioral tasks with actual K score (column 2) and predicted K score (column 3). Names of the additional behavioral tasks are listed (column 1). To investigate the relationship between predicted K scores and all of the other tasks, we correlated column 2 and column 3. We did this separately for the color and the shape tasks. The numbers in this table only reflect data from the color task. Numbers from the shape task are very similar. *P* values are parametric and uncorrected for multiple comparisons because we were interested in the overall pattern of model generalizability rather than the significance of prediction for particular behavioral measures. Further, some behavioral measures are from the same task and thus are not independent (e.g., spatial precision for different set sizes). However, we note that *p* values < .002 survive conservative Bonferroni correction for 25 comparisons.

| **Behavioral cross-correlation** |  |  |  |  |
| --- | --- | --- | --- | --- |
| **Task** | **(measured K, task)** |  | **(predicted K, task)** |  |
|  | **r** | **p** | **r** | **p** |
| Antisaccade | 0.44 | <0.0001 | 0.25 | 0.003 |
| OSPAN | 0.37 | <0.0001 | 0.05 | 0.53 |
| RSPAN | 0.36 | <0.0001 | 0.14 | 0.09 |
| Symm SPAN | 0.41 | <0.0001 | 0.14 | 0.09 |
| Average SPAN | 0.42 | <0.0001 | 0.12 | 0.15 |
| Free recall | 0.28 | 0.0008 | 0.16 | 0.06 |
| Free recall repeats | -0.09 | 0.28 | 0.08 | 0.37 |
| Paired associates | 0.29 | 0.0005 | 0.08 | 0.34 |
| Picture source | 0.31 | 0.0002 | 0.06 | 0.52 |
| Ravens | 0.26 | 0.0022 | 0.10 | 0.24 |
| Cattell | 0.39 | <0.0001 | 0.16 | 0.058 |
| Number series | 0.23 | 0.0068 | 0.12 | 0.15 |
| gF | 0.41 | <0.0001 | 0.18 | 0.038 |
| Color precision<br>(set size 1) | 0.02 | 0.84 | 0.06 | 0.50 |
| Color precision<br>(set size 6) | 0.05 | 0.59 | 0.0003 | 0.99 |
| Motion precision<br>(set size 1) | -0.31 | 0.0002 | -0.09 | 0.28 |
| Motion precision<br>(set size 6) | -0.27 | 0.0015 | -0.14 | 0.10 |
| Orientation precision<br>(set size 1) | -0.37 | <0.0001 | -0.10 | 0.25 |
| Orientation precision<br>(set size 6) | -0.44 | <0.0001 | -0.15 | 0.09 |
| Shape precision<br>(set size 1) | -0.26 | 0.0018 | -0.07 | 0.43 |
| Shape precision<br>(set size 6) | -0.09 | 0.32 | -0.04 | 0.66 |
| Spatial precision<br>(set size 1) | -0.05 | 0.57 | 0.05 | 0.53 |
| Spatial precision<br>(set size 6) | -0.30 | 0.0003 | 0.005 | 0.95 |
| Average precision from<br>all tasks (set size 1) | -0.28 | 0.0008 | -0.03 | 0.69 |
| Average precision from<br>all tasks (set size 6) | -0.39 | <0.0001 | -0.14 | 0.11 |

## DISCUSSION

Here, we demonstrate that trial-evoked EEG connectivity patterns are idiosyncratic to each person and can be used to identify individuals from a group. These patterns are unique and stable over time, just like a fingerprint. Furthermore, individual differences in these evoked EEG connectivity patterns are cognitively meaningful, predicting general fluid intelligence across individuals and working memory capacity across independent datasets. Together, these results demonstrate that individual differences in critical cognitive abilities are reflected in individuals’ unique, idiosyncratic expression of shared EEG connectivity networks. Furthermore, EEG connectivity is a generalizable and accessible approach for predicting individual differences in other abilities and behaviors from brain data.

### Predicting trait-like behavior

Previous research has used EEG to track moment-to-moment fluctuations in working memory storage. For example, multivariate pattern classification techniques have used the topography of raw EEG amplitude to decode the amount of information maintained in working memory on a single trial (Adam et al., 2020). Another EEG signal, the contralateral delay activity (CDA), scales with the number of items held in working memory. This signal’s amplitude asymptotes when working memory is full and is correlated with working memory capacity (Unsworth et al., 2015). Thus, the presence of EEG signals that track working memory storage and capacity is well established. However, no previous work has demonstrated the ability to predict working memory capacity and general fluid intelligence across completely independent individuals from different datasets using direct measures of brain function, including EEG.

Brain-based models of behavior are most theoretically informative and practically useful when they generalize to novel data. However, there are many reasons why brain-based models might not generalize. For example, internally validated model results could be driven by similarities between people at a particular site or idiosyncrasies of a particular task design or experimental context. Testing models on independent datasets (i.e., external validation) is a powerful way to reduce these and other biases (Poldrack et al., 2020; Woo et al., 2017). Although significant work has emphasized the importance of external validation (Poldrack et al., 2020), it is still relatively uncommon (Woo et al., 2017). In fact, one paper found that only 9% of the neuroscience studies that they surveyed tested models on one or more independent datasets (Woo et al., 2017). Additionally, none of these studies analyzed EEG data. Here, we externally validated our EEG connectome-based predictive models. We illustrate that our results are robust to differences in task design, experimental context, EEG system, and participant population. By externally validating our results, we provide a powerful demonstration that our models of working memory are both robust and generalizable.

Interestingly, we were able to predict general fluid intelligence using the same EEG connectivity network that predicted working memory capacity. This suggests that the variance in working memory capacity that our EEG connectivity model explains is shared with general fluid intelligence. With our cross-correlation analysis, we also found that the relationship between predicted working memory capacity and other cognitive abilities was analogous to the relationship between observed working memory capacity and other cognitive abilities. This is another example of how our model predictions capture variance in working memory that is shared with related cognitive abilities. Overall, the EEG-based connectome that we identified is not idiosyncratic to a particular working memory task. Rather, it seems to more generally track trait-like cognitive abilities, including general fluid intelligence.

### Sparse cognitive networks

Previous work in cognitive and network neuroscience has suggested that cognitive abilities, such as working memory and attention, emerge from interactions between dozens or hundreds of brain regions in complex, large-scale brain networks (Bressler & Menon, 2010; Rosenberg et al., 2017). Significant research, typically using fMRI, has utilized these networks to describe and predict behavior (Beaty et al., 2018; Finn et al., 2015; Rosenberg et al., 2016; Sripada et al., 2013; Yamashita et al., 2018). Interestingly, using EEG, we were able to predict working memory capacity from fewer than 15 functional connections and fluid intelligence from only 6 functional connections between occipital and parietal electrodes. Thus, although cognitive abilities may involve the interaction between large numbers of disparate brain regions, they can also be summarized using a relatively sparse EEG connectivity network.

### Tracking brain function

Our results suggest that common and unique signals of cognition are present in temporally precise, but spatially insensitive, EEG signals. This suggests that millisecond-by-millisecond fluctuations in neural processing are unique across individuals and cognitively meaningful. Previous work has also shown that analogous spatially sensitive signals are present in fMRI functional connectivity networks. These two methods track different neural signatures. But, when considered together, they provide additional evidence that connectivity methods, in general, track meaningful variation in brain function.

Despite the robustness of our EEG and previous fMRI functional connectivity results, there are certain obstacles that need to be addressed before we can fully accept that these methods track brain function. For example, previous work using fMRI functional connectivity has been criticized because it measures blood oxygenation, which is an indirect measure of neural activity. Due to this, critics have suggested that fMRI functional connectivity fingerprinting could be driven by individual differences in brain structure or vasculature, rather than meaningful differences in brain function (Dubois & Adolphs, 2016; Llera et al., 2019). They additionally suggest that identification and behavioral prediction could be driven by trait-like head motion, which is a challenging confound in fMRI (Nentwich et al., 2020; Siegel et al., 2017; Xifra-Porxas et al., 2020). Our EEG-based connectome models complement this work by controlling for and ruling out these potential confounds.

EEG—like MRI and all other measures of human brain activity—is also influenced by brain structure. Unlike MRI, however, EEG connectivity measures the topography of synchronous electrical brain activity, which is a direct measure of neural activity. EEG is also less influenced by head motion because EEG trials are typically shorter, and trials with head motion are removed from analyses. Therefore, these factors do not influence our ability to identify individuals and predict behavior using EEG connectivity. Nevertheless, EEG connectivity analyses come with their own unique challenges. For example, identification of individuals could be driven by differences in skull thickness across participants. Skull thickness could influence the conduction of neural signals on the scalp, leading to idiosyncratic patterns of electrical activity across people that are unrelated to brain function. We addressed this potential issue of volume conduction in the supplemental section by applying a Laplacian transformation, which reduces the adverse effects of volume conduction (Kayser & Tenke, 2015). Even when we account for potential differences in volume conduction on the scalp, we are still able to identify individuals. Overall, our results provide strong evidence that robust predictions of cognitive ability do not require the kind of dense connectivity networks that have been use in past fMRI work. EEG connectivity tracks meaningful variation in neural activity that is related to individual differences in cognition.

### Limitations and future directions

We analyzed EEG data while participants performed variants of a lateralized change detection task. Therefore, it is not clear whether predictions of working memory and general fluid intelligence rely on activity evoked during this specific task state, or whether models would generalize to predict behavior from data collected in other contexts. For example, are behaviorally relevant signals present in intrinsic EEG connectivity—that is, EEG connectivity observed at rest? Are individual differences in a cognitive process best predicted by EEG connectivity observed during a task that taxes that particular process (*c.f.,* Finn et al., 2017)? To address these questions, future research can use EEG connectivity collected during rest and different tasks to predict individual differences in other cognitive abilities, such as attentional control or long-term memory.

One complication with analyzing resting-state EEG data is that our evoked EEG connectivity measure may rely on time-locked data that is averaged over a lot of trials. This is in contrast to fMRI functional connectivity analyses, which concatenate trial data. We calculated time-locked averaged EEG data because single-trial data were noisy (see Supplemental for further detail). Resting state EEG data is typically collected continuously for a specific amount of time. It does not have specific trials, and, therefore, it is not clear how to overcome low signal-to-noise ratios with continuous resting-state EEG data. Nevertheless, it would be theoretically impactful to determine whether there are behaviorally relevant EEG networks that are present at rest.

### Conclusions

We demonstrate that a temporally precise, but sparse network of electrical activity identifies individuals and predicts cognitive abilities. The sparsity of this network suggests that dense functional networks with relatively high spatial resolution are not uniquely informative of behavior. Instead, cognitively meaningful variability in functional brain organization is also reflected in sparse, high-frequency neural signals. In sum, our analyses, which are temporally sensitive and easy and affordable to implement, provide a new arena in which we can better track and understand important individual differences in cognitive abilities.

## METHODS

Data were collected at two different study sites: the University of Oregon Eugene and the University of Chicago. Whereas data from the University of Oregon were collected as part of a single study, data from the University of Chicago were collected as multiple studies. Participants were recruited from the respective university network and from their surrounding communities. Some of these samples are described in previous publications, while others are unpublished. All data are described below.

### Subject information

#### Oregon study

Data collected at Oregon were part of one study, and the results from this dataset were previously published (Unsworth et al., 2015). All participants gave written informed consent according to procedures approved by the University of Oregon institutional review board. Participants were compensated for participation with course credit or monetary payment ($8/hr for behavior, $10/hr for EEG). Our analyses include data from 171 individuals. The numbers of participants included in our study is different than the number of participants included in the original study because we only required that participants have good EEG and change detection data, whereas their analyses required participants to have completed all of their tasks.

#### Chicago studies

Data from the University of Chicago were collected by multiple experimenters for multiple independent studies. Experiments were selected for inclusion based on whether they included a lateralized change detection task. All lateralized change detection EEG experiments that have been run by the Awh/Vogel lab at the University of Chicago task were included.

Experimental procedures were approved by The University of Chicago Institutional Review Board. All participants gave informed consent and were compensated for their participation with cash payment ($15 per hour); participants reported normal color vision and normal or corrected-to-normal visual acuity. For the current analyses, we combined all University of Chicago data into one large sample. Of note, some individuals participated in more than one University of Chicago experiments. For these individuals, behavioral data and EEG signal time-series for all trials from all of the experiments in which they participated were averaged. Across all of the Chicago studies, there were 165 unique individuals.

Chicago study #1 was previously published (Hakim, Adam, et al., 2019) and includes data from four separate experiments (28 participants in experiment 1a, 20 in experiment 1b, 20 in experiment 2a, and 29 in experiment 2b). Chicago study #2 was previously published (Hakim, Feldmann-Wüstefeld, et al., 2019) and includes data from two separate experiments (20 participants in experiment 1 and 20 participants in experiment 2). Chicago study #3 is currently in prep (Hakim et al, in prep). This study contains data from two separate experiments, however, we only analyzed data from experiment 1 (21 participants) because this was the only experiment that included a lateralized change detection task. Chicago study #4 is not published and contains data from one experiment (20 participants). Chicago study #5 is not published and contains data from two experiments (20 participants in experiment 1 and 19 participants in experiment 2). Chicago study #6 is not yet published and contains data from two experiments (25 participants in experiment 1 and 19 participants in experiment 2).

#### Task information

In all experiments, participants performed a lateralized change detection task. At the beginning of each trial, a central cue was presented to indicate which side (left or right) of the screen to pay attention to (Chicago study #6 did not have a cue on a subset of trials. See below for further information). Following the arrow cue, a memory array appeared, which consisted of a series of objects on both sides of the screen. Participants were instructed to remember the objects that were presented on the cued side of the screen, while ignoring the objects on the other side. Following the memory array, there was a blank retention interval. The exact duration of the retention interval varied across experiments. The response screen then appeared, which consisted of one object on each side of the screen. Participants had to indicate if the object on the cued side was identical to the original object that was presented at that location (Chicago study #5 had a two-alternative forced choice response. See below for further information). The exact duration and stimulus parameters varied across experiments. For published datasets, these details can be found in the original publications (Hakim, Feldmann-Wüstefeld, et al., 2019, 2019; Unsworth et al., 2015). For unpublished datasets, details are described below.

#### Chicago study #4

Stimuli were presented on a 24-in. LCD computer screen (BenQ XL2430T; 120-Hz refresh rate) on a Dell Optiplex 9020 computer. Participants were seated with their heads on a chin rest 74 cm from the screen. Each trial began with a blank inter-trial interval (750 ms), followed by a diamond cue (500 ms) indicating the relevant side of the screen (right or left). This diamond cue (maximum width = 0.65°, maximum height = 0.65°) was centered 0.65° above the fixation dot and was half green (RGB value: = 74, 183, 72) and half pink (RGB value: = 183, 73, 177). Half of the participants were instructed to attend to the green side, and the other half were instructed to attend to the pink side. After the cue, colored squares (1.1° × 1.1°) briefly appeared in each hemifield (150 ms) with a minimum of 2.10° (1.5 objects) between each square. Four colored squares appeared on each side of the screen, and then disappeared for 2000 ms. Squares could appear within a subset of the display subtending 3.1° to the left or right of fixation and 3.5° above and below fixation. Colors for the squares were selected randomly from a set of nine possible colors (RGB values: red = 255, 0, 0; green = 0, 255, 0; blue = 0, 0, 255; yellow = 255, 255, 0; magenta = 255, 0, 255; cyan = 0, 255, 255; orange = 255, 128, 0; white = 255, 255, 255; black = 1, 1, 1). Colors were chosen without replacement within each hemifield, and colors could be repeated across, but not within, hemifields. After the retention interval, the response screen then appeared. The response screen consisted of one object on each side of the screen. Participants had to report whether the color of the object on the cued side changed colors. On a subset of trials, a series of colored squares appeared on the midline during the retention interval. We did not include these trials in our analyses.

#### Chicago study #5

The experimental parameters replicate those from Chicago study #4, expect for the following changes. In Chicago study #5 experiment 1, the memory array remained on the screen throughout the delay on a subset of trials. These trials were not included in our analyses. However, because the memory array remained on the screen throughout the delay, the response screen was different than all of the other experiments. On each side of the screen, there was one object with two colors on each side of the screen (Tsubomi et al., 2013), and participants had to report which of the two presented colors the original memory item was. In Chicago study #5 experiment 2, the memory array consisted of two or four colored squares on each side of the screen, and following the memory array, the screen remained blank for 1,650 ms.

#### Chicago study #6

This data consisted of two experiments. Stimuli in all experiments were presented on a 24-in. LCD computer screen (BenQ XL2430T; 120-Hz refresh rate) on a Dell Optiplex 9020 computer. Participants were seated with their heads on a chin rest 74 cm from the screen. Each trial began with a blank inter-trial interval (1000 ms), followed by a gray cross (600 ms). The cue was either pointed to the attended location (50% of trials) or was uninformative (50% of trials). Following the cue, the memory array appeared (200 ms), which consisted of two (50% of trials) or four (50% of trials) colored target squares that appeared either to the left, right, above or below fixation. Four colored distractor circles also appeared in a position adjacent to the target squares. The display was visually balanced with four gray (RGB value: 128, 128, 128) circles across from both the target squares and the distractor circles, which matched the average luminance of the colors. Participants were told to remember the colors of the squares over the delay and ignore the circles. Squares had a side length of 0.9° visual angle and circles had a diameter of 1.0° (squares and circles covered the same area, viz., 3600 pixels). Following the memory array, the screen remained blank for 1000 ms. After this, the response screen appeared, which consisted of one colored square. Participants had to determine whether the colored square was the same color as the original colored square in that location. Colors for the target squares and distractor circles were selected randomly from a set of nine possible colors (RGB values: red = 255, 0, 0; green = 0, 255, 0; blue = 0, 0, 255; yellow = 255, 255, 0; pink = 255, 0, 255; cyan = 0, 255, 255; purple = 128, 0, 255; dark green = 4, 150, 60; orange = 255, 128, 0). Colors were chosen without replacement.

### EEG acquisition and artifact rejection

#### University of Oregon

EEG was recorded from 22 standard electrodes sites in an elastic cap (ElectroCap International, Eaton, OH) spanning the scalp, including International 10/20 sites F3, Fz, F4, T3, C3, Cz, C4, T4, P3, Pz, P4, T5, T6, O1, and O2, along with nonstandard sites OL, OR, PO3, PO4, and POz. Two additional electrodes were positioned on the left and right mastoids. All sites were recoded with a left-mastoid reference, and the data were re-referenced offline to the algebraic average of the left and right mastoids. To detect blinks, vertical electrooculogram (EOG) was recorded from an electrode mounted beneath the left eye and referenced to the left mastoid. The EEG and EOG signals were amplified with an SA Instrumentation amplifier (Fife, Scotland) with a bandpass of 0.01–80 Hz and were digitized at 250 Hz in Labview 6.1 running on a PC. Offline, data were low-pass filtered at 50 Hz to eliminate 60 Hz noise from the CRT monitor. Eye movements (>1°), blinks, blocking, drift, and muscle artifacts were detected by applying automatic criteria. This pipeline was different from the pipeline that was used in the original paper, which is why our analyses include more participants. A sliding-window step function was used to check for eye movements in the EOG channels. We used a split-half sliding-window approach (window size = 100 ms, step size = 50 ms, vertical threshold = 75 μV, horizontal threshold = 15 μV).

#### University of Chicago

EEG was recorded from 30 active Ag/AgCl electrodes (actiCHamp, Brain Products, Munich, Germany) mounted in an elastic cap positioned according to the international 10-20 system (Fp1, Fp2, F7, F8, F3, F4, Fz, FC5, FC6, FC1, FC2, C3, C4, Cz, CP5, CP6, CP1, CP2, P7, P8, P3, P4, Pz, PO7, PO8, PO3, PO4, O1, O2, Oz). Two additional electrodes were affixed with stickers to the left and right mastoids, and a ground electrode was placed in the elastic cap at position Fpz. Data were referenced online to the right mastoid. For Chicago studies #1-5, data were re-referenced offline to the algebraic average of the left and right mastoids, and incoming data were filtered (low cutoff = .01 Hz, high cutoff = 80 Hz; slope from low to high cutoff = 12 dB/octave) and recorded with a 500-Hz sampling rate. For Chicago study #6, data were re-referenced offline to the average of all electrodes, and incoming data were filtered (low cutoff=.01 Hz, high cutoff=250 Hz; slope from low to high cutoff = 12 dB/octave) and recorded with a 1000-Hz sampling rate. For all datasets, impedance values were kept below 10 kΩ. Eye movements and blinks were monitored using electrooculogram (EOG) activity and eye tracking. EOG data were collected with five passive Ag/AgCl electrodes (two vertical EOG electrodes placed above and below the right eye, two horizontal EOG electrodes placed ~1 cm from the outer canthi, and one ground electrode placed on the left cheek). Eye tracking data was collected using a desk-mounted EyeLink 1000 Plus eye-tracking camera (SR Research, Ontario, Canada) sampling at 1,000 Hz.

For Chicago studies #1-5, eye movements, blinks, blocking, drift, and muscle artifacts were first detected by applying automatic criteria. After automatic detection, trials were manually inspected to confirm that detection thresholds were working as expected. For the automatic eye movement detection pipeline, a sliding-window step function was used to check for eye movements in the horizontal EOG (HEOG) and the eye-tracking gaze coordinates. For HEOG rejection, we used a split-half sliding-window approach (window size = 100 ms, step size = 10 ms, threshold = 20 μV). HEOG rejection was only used if the eye-tracking data were bad for that trial epoch. We slid a 100-ms time window in steps of 10 ms from the beginning to the end of the trial. If the change in voltage from the first half to the second half of the window was greater than 20 μV, it was marked as an eye movement and rejected. For eye-tracking rejection, we applied a sliding-window analysis to the x-gaze coordinates and y-gaze coordinates (window size = 100 ms, step size = 10 ms, threshold = 0.5° of visual angle). We additionally used a sliding-window step function to check for blinks in the vertical EOG (window size = 80 ms, step size = 10 ms, threshold = 30 μV). We checked the eye tracking data for trial segments with missing data points (no position data are recorded when the eye is closed). We checked for drift (e.g., skin potentials) by comparing the absolute change in voltage from the first quarter of the trial to the last quarter of the trial. If the change in voltage exceeded 100 μV, the trial was rejected for drift. In addition to slow drift, we checked for sudden step-like changes in voltage with a sliding window (window size = 100 ms, step size = 10 ms, threshold = 100 μV). We excluded trials for muscle artifacts if any electrode had peak-to-peak amplitude greater than 200 μV within a 15-ms time window. We excluded trials for blocking if any electrode had at least 30 time points in any given 200-ms time window that were within 1 μ V of each other.

For Chicago study #6, eye movements, blinks, blocking, drift, and muscle artifacts were detected by applying automatic criteria only. To identify eye-related artifacts, eye-tracking data were first baselined identically to EEG data (i.e., subtraction of the mean amplitude of x and y coordinates for the time from −200 to 0 ms). Then, the Euclidian distance from the fixation cross was calculated from baselined data. Saccades were identified with a step criterion of 0.6° (comparing the mean position in the first half of a 50 ms window with the mean position in the second half of a 50-msec window; window moved in 20 ms steps). Drifts were identified by eyetracking data indicating a distance from the fixation of >1°. Both eyes had to indicate an eye-related artifact for a trial to be excluded from analysis. In addition, trials in which any EEG channel showed a voltage of more than 100 μV or less than −100 μV were rejected.

#### Behavioral analysis

Working memory capacity (*K score*) was used to measure task performance (Cowan, 2001; Pashler, 1988). The formula that we used was *K* = *S*(*H* – *F*), where *K* is working memory capacity, *S* is the size of the array, *H* is the observed hit rate, and *F* is the false alarm rate.

#### EEG connectivity analyses

We first tested our analyses within the Oregon dataset, whenever this was possible. We then replication these analyses and externally validated predictive models in the compiled Chicago study.

##### Calculation of EEG functional connectivity

The Chicago and Oregon datasets had different numbers of electrodes. Therefore, in our analyses we only included the electrodes that overlapped between the two datasets. This left us with a total of 17 electrodes: F3, Fz, F4, C3, Cz, C4, P7, P3, Pz, P4, P8, PO8, PO4, PO3, PO7, O1, and O2. For each participant, we averaged the raw amplitude at each of these 17 electrodes across all trials from timepoints 0 to 1000 ms. Timepoint zero corresponded to the onset of the memory array, and timepoint 1000 was the end of the shortest retention interval (1000 ms). We analyzed data from the same time window for all experiments to match the number of timepoints included in the analysis across experiments. We computed the Pearson correlation of this trial-averaged EEG activity for all pairwise electrodes for each participant separately. For each participant, this resulted in a 17 x 17 matrix of the correlation between the time course of each electrode to each other electrode. We Fisher z-transformed correlation coefficients and submitted the resulting functional connectivity matrices to the analyses described below.

##### Functional connectivity fingerprinting

This analysis investigates whether individuals’ EEG connectivity patterns are unique and stable enough to distinguish them from a group. These methods were adopted from a previous paper that investigated functional connectome fingerprinting in fMRI (Finn et al., 2015).

To identify individuals in the Oregon dataset, we compared participants’ vectorized shape-task and color-task EEG connectivity matrices. Specifically, for each individual, we correlated their “target” color-task EEG connectivity vector from with a “database” of all shape-task EEG connectivity vectors. This individual was considered accurately identified if the maximum correlation was with their own shape-task data. We repeated this analysis using shape-task vectors as the “targets” and color-task vectors as the “database”. Finally, we characterized functional connectome fingerprinting accuracy as the number of correct identifications divided by the number of participants*2.

For the Chicago study, we analyzed data from the subset of 45 individuals who participated in more than one experiment. For each person in this sample, we first calculated and vectorized an EEG functional connectivity matrix from each experiment in which they participated separately. Next, we selected one of these vectors to serve as the “target”. We compared this target to a database, which included 19 individuals’ connectivity vectors from another task in which the target individual also participated (including the target individual). We chose 19 because this was the minimum number of participants in any individual study, and we wanted to equate chance levels across the experiments. To predict subject identity, we computed the similarity between the target connectivity vector and each database vector, and the predicted identity was that with the maximal similarity score. Similarity was defined as the Pearson correlation between the vectors.

To ask whether functional connections that predict behavior contribute to identification more than expected by chance, we ran these analyses including only edges that significantly predicted behavior. To determine whether these results were due to down sampling the number of edges included in the connectivity matrix, we compared the results from the significant edge analysis to an analysis with the same number of randomly selected edges.

To assess the statistical significance of identification accuracy, we performed nonparametric permutation tests. We ran the same analysis as above, except we shuffled the subject labels in each iteration, so that they would randomly align with the EEG connectivity data. This shuffling of labels was repeated 10,000 times.

##### Connectome-based predictive modeling

Prediction methods were adopted from previous papers that used fMRI functional connectivity to predict behavior (Rosenberg et al., 2016; Shen et al., 2017). We first separated data into training and testing sets. For internally validated models, we used 5-fold cross-validation and permutation testing to determine whether the model significantly predicted behavior. For external validation analyses, we trained models using data collected in Oregon and tested them using data collected in Chicago and vice versa.

To identify the edges that were significantly related to behavior in the training sample, we correlated each edge in the connectivity matrix to K score using Spearman’s correlation. Then, we selected the top 10% of edges that most strongly predicted behavior in the positive and negative directions. (See supplemental section for more details on using different thresholds to select features.) Using these features, we calculated single-subject summary values for all training subjects. To do this, we summed the edge-strength values for each individual for both the positive and negative edges separately, and then we took the difference between them. We trained a linear model with these summary features and K scores from the training set to predict behavior from EEG connectivity. We then used this model to predict K score in the testing set. To do this, we calculated summary features in test set participants, and input these summary scores into the model defined in the training sample to generate predictions for each test set subject’s K score. Results are reported as p-values, r-values, and mean square error (mse).

##### Predictive network overlap

We calculated whether the predictive edges from the Oregon and Chicago models significantly overlapped. We determined significance of network overlap using the hypergeometric cumulative density function in Matlab (Rosenberg et al., 2016). The formula was as follows: *p*=1-hygecdf(*x, M, K, N*), where *x* is the number of overlapping edges, *M* is the total number of edges in the matrix, *K* is the number of edges in the Oregon network, and *N* is the number of edges in the Chicago network.

##### Characterizing the relationship between predicted working memory and other cognitive abilities

Finally, we asked whether the models that we used to predict working memory capacity across individuals were related to other cognitive abilities. We ran these analyses only with the Oregon dataset because these participants completed behavioral tasks assessing a range of cognitive abilities in addition to the change detection tasks used to measure K. These analyses included the 138 participants who completed all cognitive tasks.

We were specifically interested in the relationship between EEG connectivity and gF. To investigate this relationship, for each subject in the Oregon dataset, we calculated the strength in the working memory network defined in the Chicago sample. Then, using 5-fold crossvalidation, we defined a linear model relating network strength to gF. We then used this model to predict the left-out set of participants’ gF scores. Previous research has shown that gF and K score are highly correlated (Conway et al., 2002; Unsworth et al., 2015). Given this relationship, we calculated a theoretical ceiling of these models’ performance as the correlation between gF and K scores.

As an exploratory analysis, we also investigated the relationship between predicted K scores and performance on all other behavioral tasks (**Table 1**). To do this, we computed the Pearson correlation between 1) observed K score and all of the other tasks (**Table 1**, column 2) and 2) predicted K score and all of the other tasks (**Table 1**, column 3). Then, we correlated these two columns of data to determine whether there was a significant relationship between performance on all tasks and the measured and predicted K scores. We did this separately for the color and the shape tasks.

## ACKNOWLEDGMENTS

Research was supported by NIMH grant ROIMH087214 and Office of Naval Research grant N00014-12-1-0972. We thank Tobias Feldmann-Wüstefeld for allowing us to use his dataset in our analyses and Kirsten C.S. Adam for organizing and pre-processing the Oregon site data.

## AUTHOR CONTRIBUTIONS

NH and MDR conceived of the study. NH ran the analyses with support and contributions from MDR. NH and MDR wrote the paper with contributions from EA and EKV.

