## Supplemental Materials for "Predicting cognitive abilities across individuals using sparse EEG connectivity"

#### Model availability.

All of our data and analysis scripts will be available online upon publication of this manuscript. We have additionally created and uploaded a model that is trained on the combination of the Oregon-site and Chicago-site datasets. This model can be used for future research that is interested in investigating EEG connectivity.

#### Model building.

When we were originally building our model, we ran all of our analyses within the Oregon-site data because it was the largest sample available and all participants performed the same two lateralized change detection tasks. Once the model was finalized, we applied it to the Chicago-site data. We then built a model on the Chicago sample and externally validated it using the Oregon data. Thus, the model-fitting analyses described below (*Concatenating versus averaging trials*, *Electrode organization*, and *Fingerprinting and behavioral predictions with trial-evoked amplitude*) were only run on the Oregon-site data.

#### Concatenating versus averaging trials.

Previous fMRI connectome-based models have concatenated data across trials and then correlated these concatenated time series across all pairwise electrodes. Our EEG data, however, were very noisy when we concatenated the data across trials and did not significantly predict behavior in the Oregon-site data: median $r$=0.02, $p$=0.39, *mse*=0.27, *Cohen's d*=0.26; **Supplemental figure 1**. Therefore, instead of concatenating trials, we averaged the data across trials and then correlated these averaged time series across all pairwise electrodes. This resulted in a trial-evoked EEG-based connectivity signal. Trial-evoked signals eliminate any neural activity that was not consistent across trials. As noted in the discussion section, future research should investigate ways to track other EEG signals that are not trial-evoked.

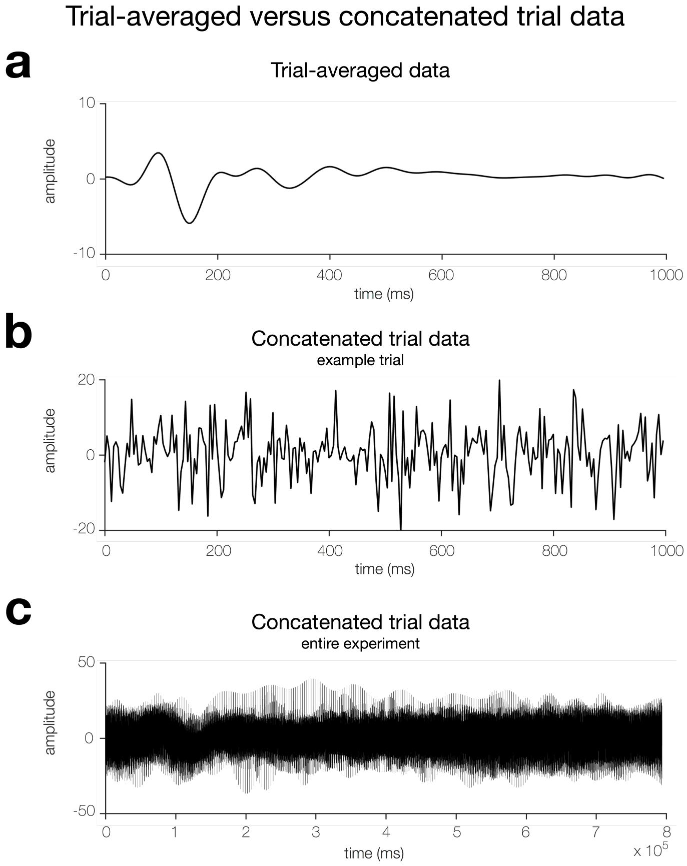


Supplemental figure 1 | Example of trial-averaged versus concatenated trial data. All of the figures are data from participant #1, electrode PO8. They are representative of the sample at large. **a** Example of the type of data that we used for our connectivity analyses. This data is averaged over all trials for each of the 17 electrodes. This figure contains data only from PO8. **b** Example of what the data look like if we concatenate all trials, instead of averaging across trials. This figure is the time-course from a single example trial, so that the time courses for (**a)** and (**b**) can be directly compared. **c** Example of what the data look like if we concatenate all trials, instead of averaging across trials. This figure shows the time course of electrode PO8 across the entire experiment.

**Electrode organization**.

We analyzed EEG data that was collected while participants performed a lateralized change detection task. In this type of task, participants have to attend and maintain information on either the left or right side of the screen on any given trial. This results in lateralized neural activity that is contralateral to the remembered items (Vogel & Machizawa, 2004). For example, if stimuli are presented on the left side of the screen, this lateralized neural activity would be present in electrodes O2, PO8, PO4, etc. Whereas if stimuli were presented on the right side of the screen, it would be present in electrodes O1, PO7, PO3 We accounted for this lateralized neural activity by aligning neural activity based on which side of the screen participants were attending on each trial. For example, activity for electrode number 10 in the functional connectivity matrix included data from PO8 on "attend-left" trials and data from PO7 on "attend-right" trials. Central electrodes (Fz, Cz, and Pz) were unaffected by this organization. This method of alignment is similar to how CDA analyses account for the contralateral organization of the visual system (Vogel & Machizawa, 2004). When we did not flip the electrodes in this way, we were unable to predict working memory capacity in the Oregon dataset: $r$=0.02, $p$=0.29, *mse*=0.27, *Cohen's d*=0.25.

**Fingerprinting and behavioral predictions with trial-evoked amplitude**

We sought to determine whether trial-evoked EEG amplitude could identify individuals and predict behavior across individuals. If this is the case, this would suggest that we do not need to account for the relationship between electrodes in order to identify individuals and predict cognitive abilities. Therefore, we ran the fingerprinting analysis and built a model to predict behavior within the Oregon-site data using vectors of trial-averaged amplitude at each electrode. These analyses were identical to the ones above, however, we used trial-averaged amplitude, instead of EEG functional connectivity. This analysis revealed that we were still able to identify individuals using trial-evoked amplitude: color: shape task accuracy = 51%, $p<0.0001$; shape: color task accuracy = 51%, $p<0.0001$ (chance = 1/171, or .58%). However, identification accuracy was lower than when we used EEG functional connectivity (color: shape task accuracy = 82%, $p<0.0001$; shape: color task accuracy = 81%, $p<0.0001$). Additionally, we were not able to significantly predict behavior across individuals using amplitude: $r=0.02$, $p=0.43$, $mse=0.26$, *Cohen's d*=0.17. These results suggest that the ability to identify individuals and predict cognitive abilities across individuals relies on the relationship between different electrodes on the scalp.

**Laplacian transformed analyses**.

Differences in skull thickness across participants could overestimate the ability of EEG connectivity to identify individuals from a group because skull thickness influences the conduction of electrical brain activity on the scalp. To account for potential differences in volume conduction, we applied a Laplacian transformation to our data and then ran the identification analyses. A Laplacian transformation is a spatial filter that can increase topographical specificity by accounting for electrical activity at neighboring electrodes (Vidal et al., 2015). By accounting for nearby electrical activity, the Laplacian transformation can account for how much a signal has spread across the scalp. We found that even when we accounted for potential differences in volume conduction, we were still able to identify individuals with high accuracy in both the Oregon-site data: color: shape task accuracy = 77.78%, $p<0.001$; shape: color task accuracy = 80.12%, $p<0.001$ (chance = 1/171, or .58%), and the Chicago-site data: task A:B *accuracy*=37.62%, $p<0.001$; task B:A *accuracy*=36.00%, $p<0.001$ (chance = 1/19, or 5.26%). This aligns with previous research that has found that differences in brain activity contribute much more variance to surface EEG than do variations in skull thickness (Hagemann et al., 2008).

**Number of predictive edges**.

The number of total edges in our EEG connectivity analyses is less than 1% of the number of edges that have been included in analogous fMRI connectome-based models. Given this large difference in the total number of edges between these two methods, we did not have strong a priori predictions about what percentage of the predictive edges we should include in our EEG connectome-based models. Previous fMRI connectome-based predictive modeling research identified predictive networks consisting of fewer than approximately 10% of available functional connections (Finn et al., 2015; Rosenberg et al., 2018). Based on this, our models included the top 5% of positive and top 5% of negative edges (10% total). Nevertheless, we were interested in investigating how the number of edges influenced model performance and the robustness of model predictions to feature selection threshold. Therefore, as an exploratory analysis, we systematically varied the number of predictive edges that were included in our externally validated EEG connectome-based models; **Supplemental figure 2**.

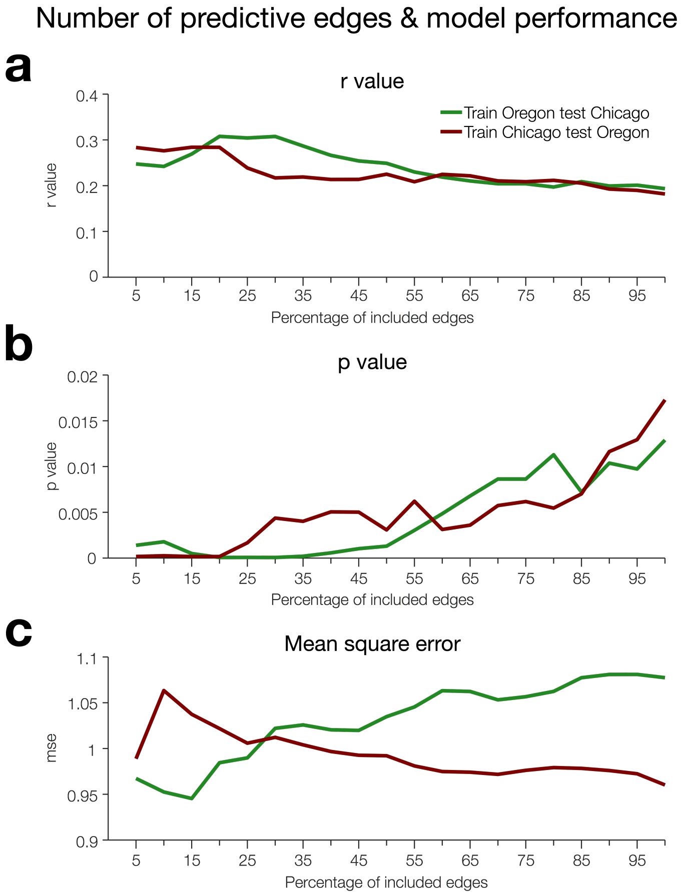


Supplemental figure 2 | Model performance based on the number of predictive edges used to calculate EEG summary features. *R*-values (**a**), *p*-values (**b**), and mean square error (**c**) associated with externally validated models when we included a certain percentage of the predictive edges to calculate summary features. The percentage of included edges (5% to 100% in steps of 5) is on the x-axis.

### How much data do we need to get robust predictions?

Our models are the first of their kind to predict cognitive abilities across completely independent datasets using evoked-EEG connectivity. Given the novelty of these analyses, we sought to determine how many testing participants should be included in future research that investigates similar questions with this new method. Given that our fully trained EEG connectome-based models are available online (see *Model availability* for detail), we sought to determine how the number of testing participants influences model performance. To determine this, we down sampled the number of participants included in the testing set of both of our externally validated models; **Supplemental figure 3**. We iterated this down sampling 10,000 times. Based on these results, we suggest that a minimum of approximately 100 testing participants should be included in future analyses of this kind.

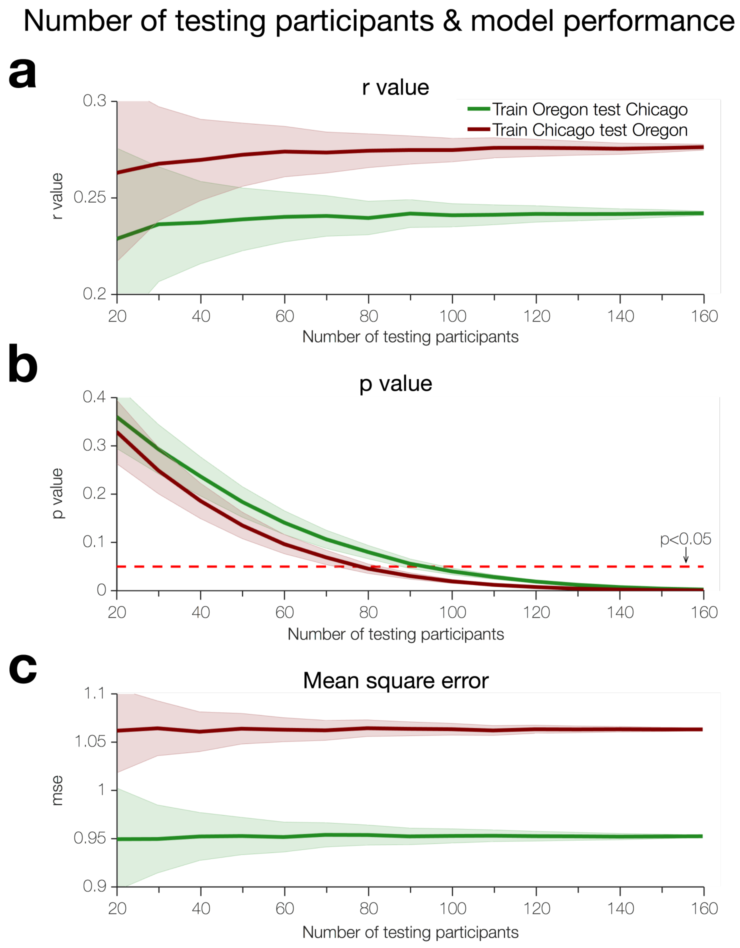


Supplemental figure 3 | Number of testing participants and model performance. *R*-values (**a**), *p*-values (**b**), and mean square error (**c**) associated with externally validated models when we included a given number of testing participants. The number of testing participants (20 to 160 in steps of 10) is on the x-axis.

**Relationship between number of trials and working memory capacity**.

The Chicago-site data is a compilation of 12 different experiments, and a given participant could have completed any number of these studies. There is some concern that participants who completed multiple studies could be different from participants who only completed one study. One possibility is that participants with higher working memory capacity may be more compliant test subjects and, thus, could have completed more studies than participants with lower working memory capacities. If this is the case, then our EEG connectivity matrices would include more trials for higher than for lower working memory capacity participants. To investigate whether this was the case, we correlated working memory capacity with the number of trials included in our analyses. We found that there was not a significant relationship between working memory capacity and number of trials ($r$=0.13, $p$=0.10, $mse$=1.65e6). Therefore, our ability to predict working memory capacity across individuals in the Chicago-site dataset was not contingent on differences in the number of trials between high and low working memory capacity individuals. This is not an issue in the Oregon-site data because all participants completed the same number of trials.
